# Do we need demographic data to forecast plant population dynamics?

**DOI:** 10.1101/025742

**Authors:** Andrew T. Tredennick, Mevin B. Hooten, Peter B. Adler

## Abstract

1. Rapid environmental change has generated growing interest in forecasts of future population trajectories. Traditional population models built with detailed demographic observations from one study site can address the impacts of environmental change at particular locations, but are difficult to scale up to the landscape and regional scales relevant to management decisions. An alternative is to build models using population-level data that are much easier to collect over broad spatial scales than individual-level data. However, it is unknown whether models built using population-level data adequately capture the effects of density-dependence and environmental forcing that are necessary to generate skillful forecasts.
2. Here, we test the consequences of aggregating individual 29 responses when forecasting the population states (percent cover) and trajectories of four perennial grass species in a semi-arid grassland in Montana, USA. We parameterized two population models for each species, one based on individual-level data (survival, growth and recruitment) and one on population-level data (percent cover), and compared their forecasting accuracy and forecast horizons with and without the inclusion of climate covariates. For both models, we used Bayesian ridge regression to weight the influence of climate covariates for optimal prediction.
3. In the absence of climate effects, we found no significant difference between the forecast accuracy of models based on individual-level data and models based on population-level data. Climate effects were weak, but increased forecast accuracy for two species. Increases in accuracy with climate covariates were similar between model types.
4. In our case study, percent cover models generated forecasts as accurate as those from a demographic model. For the goal of forecasting, models based on aggregated individual level data may offer a practical alternative to data-intensive demographic models. Long time series of percent cover data already exist for many plant species. Modelers should exploit these data to predict the impacts of environmental change.

## Introduction

Perhaps the greatest challenge for ecology in the 21st century is to forecast the impacts of environmental change (Clark et al. 2001, Petchey et al. 2015). Forecasts require sophisticated modeling approaches that fully account for uncertainty and variability in both ecological process and model parameters (Luo et al. 2011, but see Perretti et al. 2013). The increasing statistical sophistication of population models (Rees and Ellner 2009) makes them promising tools for predicting the impacts of environmental change on species persistence and abundance. But reconciling the scales at which population models are parameterized with the scales at which environmental changes play out remains a challenge (Clark et al. 2010, 2012, Freckleton et al. 2011, Queenborough et al. 2011). Most population models are built using demographic data from a single study site because tracking the fates of individuals is so difficult. The resulting models cannot be applied to the landscape and regional scales relevant to decision-making without information about how the estimated parameters respond to spatial variation in biotic and abiotic drivers (Sæther et al. 2007). The limited spatial extent of individual-level demographic datasets constrains our ability to use population models to address applied questions about the consequences of climate change.

Aggregate measures of population status, rather than individual performance, offer an intriguing alternative for modeling populations (Clark and Bjørnstad 2004, Freckleton et al. 2011). Population-level data cannot provide inference about demographic mechanisms, but might be sufficient for modeling future population states, especially because population-level data, such as plant percent cover, are feasible to collect across broad spatial extents (e.g., Queenborough et al. 2011). The choice between individual and population-level data involves a difficult trade-off: while individual-level data are necessary for mechanistic models, population-level data enable models that can be applied over greater spatial and temporal extents. An open question is how much forecasting skill is lost when we build models based on population rather than individual-level data.

To date, most empirical population modelers have relied on individual-level data, with few attempts to capitalize on population-level measures. An important exception was an effort by Taylor and Hastings (2004) to model the population growth rate of an invasive species to identify the best strategies for invasion control. They used a “density-structured” model where the state variable is a discrete density state rather than a continuous density measure. Such models do not require individual-level demographic data and can adequately describe population dynamics. Building on Taylor and Hastings (2004), Freckleton et al. (2011) showed that density-structured models compare well to continuous models in theory, and Queenborough et al. (2011) provide empirical evidence that density-structured models are capable of reproducing population dynamics at landscape spatial scales (also see Mieszkowska et al. 2013), even if some precision is lost when compared to fully continuous models. However, previous tests of density-structured population models have yet to assess their ability to forecast out-of-sample observations, and they have not included environmental covariates, which are necessary to forecast population responses to climate change.

Addressing climate change questions with models fit to population-level data is potentially problematic. Population growth (or decline) is the outcome of demographic processes such as survival, growth, and recruitment that occur at the level of individual plants. Climate can affect each demographic process in unique, potentially opposing, ways (Dalgleish et al. 2011). These unique climate responses may be difficult to resolve in statistical models based on population-level data where demographic processes are not identifiable. Futhermore, models based on aggregated data may reflect short-term effects of one vital rate more than others whose importance may only emerge over the long-term. For example, a one-year change in a plant species’ cover or biomass might reflect growth or shrinkage of the largest individuals, whereas the long-term trajectory of the population might be more influenced by recruitment. The same is true for density dependence: intraspecific density depedence may act most strongly on vital rates, like recruitment, that are difficult to identify from population-level data. If density dependence and/or important climate effects are missed because of the aggregation inherent in population-level data, then population models built with such data will make uninformative or unreliable forecasts.

We compared the forecasting skill (accuracy and precision) of statistical and population models based on aggregated, population-level data with the skill of models based on individual-level data. We used a demographic dataset that tracks the fates of individual plants from four species over 14 years to build two kinds of single-species population models, traditional models using individual growth, survival, and recruitment data and alternative models based on population-level (basal cover) data. We simulated from the models to answer two questions motivated by the fact that the effects of intraspecific competition (density dependence) and interannual weather variability act at the level of the individual (Clark et al. 2011). First, can population models fit using aggregated individual-level data (percent cover) adequately capture density dependence to produce forecasts as skillful as those from models fit to demographic data? Second, can population models fit using aggregated data adequately capture the influence of climate on population growth and, in turn, produce forecasts as skillful as those from models fit to demographic data?

## Materials and Methods

### Overview of analysis

We used two types of data: individual-level data and percent cover data. Using the individual-level data, we fit three vital rate regressions (survival, growth, and rectruitment) to build an Integral Projection Model (IPM) for simulating the plant populations. Using the percent cover data we fit a simple, Gompertz population growth model, which we refer to as a quadrat-based model (QBM). For both model types (IPM and QBM), we fit and simulate versions of the model with and without climate covariates. We used Bayesian ridge regression to weight the importance of each climate covariate. We then performed cross-validation to assess each model’s ability to predict out-of-sample data. We compared the forecast accuracy (*ρ*, correlation between observations and predictions) and mean absolute error (MAE) between the IPM and the QBM to test our expectation that the IPM should outperform the QBM. Lastly, we use in-sample forecasts to quantify each model’s forecast horizon (Petchey et al. 2015).

### Study site and data

Our demographic data were obtained from a northern mixed grass prairie at the Fort Keogh Livestock and Range Research Laboratory near Miles City, Montana, USA (46° 19’ N, 105° 48° W). The dataset is available on Ecological Archives^1^ (Anderson et al. 2011), and interested readers should refer to the metadata for a complete description. The site is 800 m above sea level and mean annual precipitation (1878–2009) is 334 mm, with most annual precipitation falling from April through September. The community is grass-dominated, and we focused on the four most abundant grass species: *Bouteloua gracilis* (BOGR), *Hesperostipa comata* (HECO), *Pascopyrum smithii* (PASM), and *Poa secunda* (POSE) (Fig. 1 and Table 1). *B. gracilis* is a warm-season perennial grass, whereas *H. comata, P. smithii*, and *Poa secunda* are cool-season perennial grasses. The growing season begins in early spring (typically in April) and lasts through midsummer (typically in June).

**Table 1.** Description of data. The observations span 13 year-to-year transitions.

| Species | Vital Rate Model | Num. Obs. | Num. Quadrats |
| --- | --- | --- | --- |
| *B. gracilis* | Growth | 5670 | 29 |
|  | Survival | 10102 | 33 |
|  | Recruitment | 304 | 33 |
|  | Percent cover | 281 | 29 |
| *H. comata* | Growth | 1990 | 16 |
|  | Survival | 3257 | 18 |
|  | Recruitment | 304 | 18 |
|  | Percent cover | 171 | 17 |
| *P. smithii* | Growth | 8052 | 19 |
|  | Survival | 11344 | 19 |
|  | Recruitment | 304 | 19 |
|  | Percent cover | 217 | 19 |
| *Poa secunda* | Growth | 3018 | 18 |
|  | Survival | 4650 | 18 |
|  | Recruitment | 304 | 18 |
|  | Percent cover | 197 | 18 |

**Figure 1.**
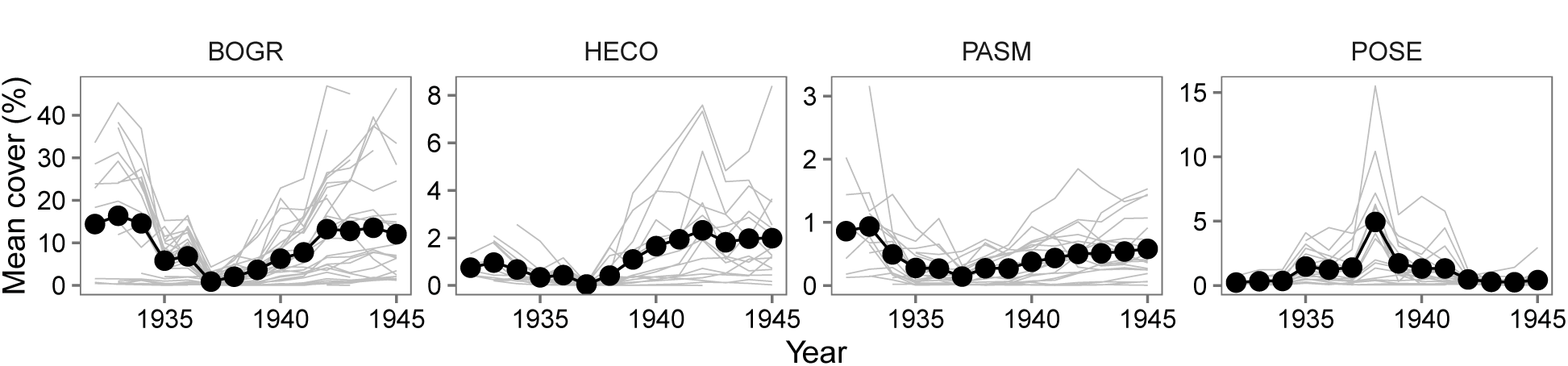
Time series of average percent cover over all quadrats for our four focal species: *Bouteloua gracilis* (BOGR), *Hesperostipa comata* (HECO), *Pascopyrum smithii* (PASM), and *Poa secunda* (POSE). Light grey lines show trajectories of individual quadrats. Note the different y-axis scales across panels. See Table 1 for sample size information.

From 1932 to 1945, individual plants were identified and mapped annually in 44 1-m^2^ quadrats using a pantograph. The quadrats were distributed among six pastures, each assigned a grazing treatment of light (1.24 ha/animal unit month), moderate (0.92 ha/aum), and heavy (0.76 ha/aum) stocking rates (two pastures per treatment). In this analysis, we accounted for potential differences among the grazing treatments, but do not focus on grazing×climate interactions. The annual maps of the quadrats were digitized and the fates of individual plants tracked and extracted using a computer program (Lauenroth and Adler 2008, Chu et al. 2014). The permanent quadrats have not been relocated, but their distribution in six different pastures implies that the data represent a broad spatial distribution for the study area. Daily climate data are available for the duration of the data collection period (1932 – 1945) from the Miles City airport, Wiley Field, 9 km from the study site.

We modeled each grass population based on two levels of data: individual and quadrat. The individual data are the “raw” data. For the quadrat-level data, we summed individual basal cover for each quadrat by species. This is equivalent to a near-perfect census of quadrat percent cover because measurement error at the individual-level is small (Chu and Adler 2015). Based on these two datasets of 13 year-to-year transitions, we can compare population models built using individual-level data and aggregated, quadrat-level data. At the individual level, we explicitly model three vital rates: growth, survival, and recruitment. At the quadrat level, we model population growth as change in percent cover of quadrats with non-zero cover in year *t* and in year *t-1*, ignoring within-quadrat extirpation and colonization events because they are very rare in our time series (*N* = 16 and *N* = 13, respectively, across all species). Sample sizes for each species and vital rate model are shown in Table 1.

All R code and data necessary to reproduce our analysis is archived on GitHub as release v1.0^2^(http://github.com/atredennick/MicroMesoForecast/releases). We have also deposited the v1.0 release on Dryad *(link here after acceptance*).

### Statistical models of vital rates

At both levels of inference (individual and quadrat), the building blocks of our population models are vital rate regressions. For individual-level data, we fit regressions for survival, growth, and recruitment for each species. At the quadrat-level, we fit a single regression model for population growth. We describe the statistical models separately because they required different approaches. For both model types, we fit vital rate models with and without climate covariates. Models with climate effects contain five climate covariates that we chose *a priori* based on previous model selection efforts using these data (Chu et al. 2016) and expert advice (Lance Vermeire, *personal communication*): “water year” precipitation at *t*-2 (lagppt); April through June precipitation at *t*-1 and *t* (ppt1 and ppt2, respectively) and April through June temperature at *t*-1 and *t* (TmeanSpr1 and TmeanSpr2, respectively), where *t*-1 to *t* is the transition of interest. We also include interactions among same-year climate covariates (e.g., ppt1 × TmeansSpr1), resulting in a total of seven climate covariates.

We fit all models using a hierarchical Bayesian approach. In the following description, we focus on the main process and the model likelihood (full model descriptions are in the Supporting Information). For the likelihood models, y^*X*^ is always the relevant vector of observations for vital rate *X* (*X* = *S, G, R*, or *P* for survival, growth, recruitment, or population growth). For example, y^*S*^ is a vector of 0s and 1s indicating whether a genet survives from *t* to *t+1*, or not, for all observation years and quadrats. All model parameters are species-specific, but we omit subscripts for species in model descriptions below to reduce visual clutter. For brevity, we only describe models with climate covariates included, but models without climate covariates are simply the models described below with the climate effects removed.

#### Vital rate models at the individual level

We used logistic regression to model the probability that genet *i* in quadrat *q* survives from time *t* to *t*+1 (*s*_*i,q*_,_*t*_):

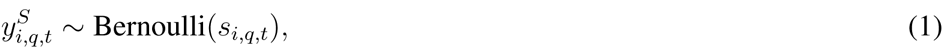

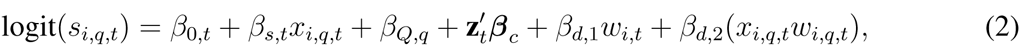

where *x*_*i,q,t*_ is the log of genet *i* basal area at time *t, β*_0*,t*_ is a year specific intercept, *β*_*Q,q*_ is the random effect of the *q*th quadrat to account for spatial location, *β*_*s,t*_ is the year-specific slope parameter for size, **z** is a vector of *p* climate covariates specific to year *t, β*_*c*_ is a vector of fixed climate effects of length *p*, *β*_*d*,1_ is the effect of intraspecific crowding experienced by the focal genet at time *t* (*w*_*i,q,t*_), and *β*_*d*,2_ is a size by crowding (*x*_*i,q,t*_*w*_*i,q,t*_) interaction effect.

We follow the approach of Chu and Adler (2015) to estimate crowding, assuming that the crowding experienced by a focal genet depends on distance to each neighbor genet and the neighbor’s size, *u*:

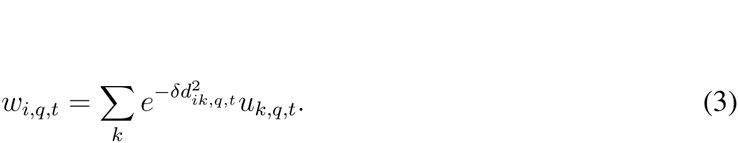

In equation 3, *w*_*i,q,t*_ is the crowding that genet *i* in year *t* experiences from *k* conspecific neighbors (*u*_*k,q,t*_) in quadrat *q*. The spatial scale over which conspecific neighbors exert influence on any genet is determined by *δ*. The function is applied for all *k* conspecific genets that neighbor the focal genet at time *t*, and *d*_*ik,q,t*_ is the distance between genet *i* and conspecific genet *k* in quadrat *q*. We use regression-specific (survival and growth) *δ* values estimated by Chu and Adler (2015).

We modeled growth as a Gaussian process describing log genet size 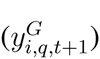 at time *t* + 1 in quadrat *q* as a function of log size at time *t* and climate covariates:

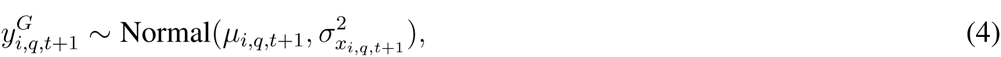

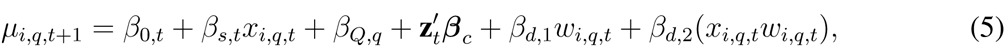

where *µ*_*i,q,t+*1_ is log of genet is predicted size at time *t+1*, and all other parameters are as described for the survival regression. We capture non-constant error variance in growth by modeling the variance in the growth regression 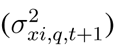 as a nonlinear function of predicted genet size:

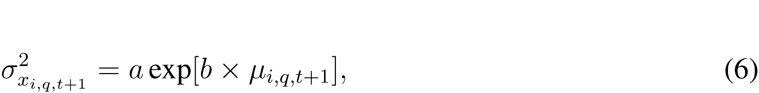

where *µ*_*i,q,t+*1_ is log of predicted genet size predicted from the growth regression (Eq. 4), and *a* and *b* are constants.

Our data allows us to track new recruits, but we cannot assign a specific parent to new genets. Therefore, we model recruitment at the quadrat level. We assume the number of individuals, 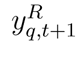, recruiting at time *t* + 1 in quadrat *q* follows a negative binomial distribution:

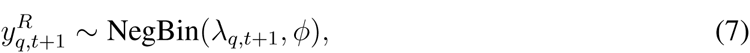

where λ is the mean intensity and *φ* is the size parameter. We define λ as a function of quadrat composition and climate in the previous year:

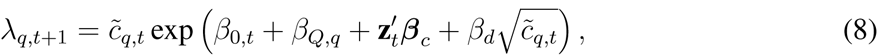

where 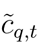 is effective cover (cm^2^) of the focal species in quadrat *q* at time *t*, and all other terms are as in the survival and growth regressions. Effective cover is a mixture of observed cover (*c*) in the focal quadrat (*q*) and the mean cover across the entire group 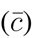 of *Q* quadrats in which *q* is located:

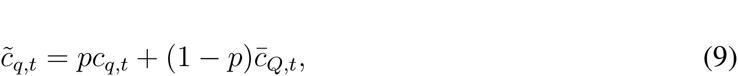

where *p* is a mixing fraction between 0 and 1 that is estimated when fitting the model.

#### Population model at the quadrat level

The statistical approach used to model aggregated data depends on the type of data collected. We have percent cover data, which can easily be transformed to proportion data in our case because plant areas were scaled by plot area. An obvious choice for fitting a linear model to proportion data is beta regression because the support of the beta distribution is (0,1), which does not include true zeros or ones. However, when we used fitted model parameters from a beta regression in a quadrat-based population model, the simulated population tended toward 100% cover for all species. We therefore chose a modeling approach based on a truncated log-normal likelihood. The model for quadrat cover change from time *t* to *t* + 1 is

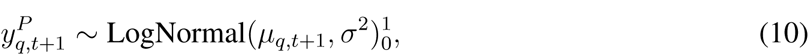

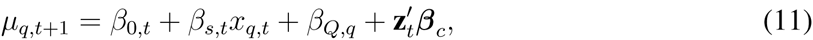

where *µ*_*q,t*+1_ is the log of proportional cover in quadrat *q* at time *t*+1, and all other parameters are as in the individual-level growth model (Eq. 4) except that *x* now represents log of proportional cover. The log normal likelihood includes a truncation (subscript 0, superscript 1) to ensure that predicted values do not exceed 100% cover.

### Model fitting and statistical regularization

#### Model fitting

Our Bayesian approach to fitting the vital rate models required choosing appropriate priors for unknown parameters and deciding which, if any, of those priors should be hierarchical. For each species, we fit yearly size effects and yearly intercepts hierarchically, where year-specific coefficients were modeled with global distributions representing the mean size effect and intercept. Quadrat random effects were also fit hierarchically, with quadrat offsets modeled using distributions with mean zero and a shared variance term (independent Gaussian priors). Climate effects were modeled as independent covariates whose prior distributions were optimized for prediction using statistical regularization (see **Statistical regularization: Bayesian ridge regression** below).

All of our analyses (model fitting and simulating) were conducted in R (R Core Team 2013). We used the ‘No-U-Turn’ Hamiltonian Monte Carlo sampler in Stan (Stan Development Team 2014a) to sample from the posterior distribution of model parameters using the package rstan (Stan Development Team 2014b). We obtained samples from the posterior distribution for all model parameters from three parallel MCMC chains run for 1,000 iterations after discarding an initial 1,000 iterations. Such short MCMC chains are possible because the Stan sampler reduces the number of iterations needed to achieve convergence. We assessed convergence visually and checked that scale reduction factors for all parameters were less than 1.1. For the purposes of including parameter uncertainty in our population models, we retained the final 1,000 iterations from each of the three MCMC chains to be used as randomly drawn values during population simulation. We report the posterior mean, standard deviation, and 95% Bayesian Credible Intervals for every parameter of each model for each species in the Supporting Information (Tables S5-S20).

#### Statistical regularization: Bayesian ridge regression

For models with climate covariates, our objective is to model the response of our focal grass species to interannual variation in climate, even if those responses are weak. Therefore, we avoid selecting among models with all possible combinations of climate covariates, and instead use Bayesian ridge regression to regulate, or constrain, the posterior distributions of each climate covariate (Gerber et al. 2015, Hooten and Hobbs 2015). Ridge regression is a specific application of statistical regularization that seeks to optimize model generality by trading off bias and variance. As the name implies, statistical regularization involves the use of a regulator that constrains an optimization. The natural regulator in a Bayesian application is the prior on the coefficients of interest. Each of our statistical models includes the effects of climate covariates via the term 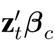 with prior *β*_*c*_ ~ Normal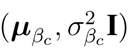. Because we standardized all climate covariates to have mean zero and variance one, we set ***µ***_*βc*_ = 0 and let 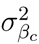 serve as the regulator that shrinks covariate effects toward zero – the smaller the prior variance, the more the posteriors of *β*_*c*_ are shrunk toward zero, and the stronger the penalty (Hooten and Hobbs 2015).

To find the optimal penalty (i.e., optimal value of the hyperparameter 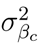), we fit each statistical model with a range of values for 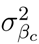 and compared predictive scores from leave-one-year-out cross-validation. We performed the grid search over 24 values of 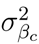, ranging from 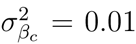 to 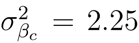. For each statistical model and each species, we fit 13 × 24 = 312 iterations of the model fitting algorithm to search 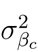 for the optimal value (13 years to leave out for crossvalidation and 24 values of 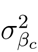) – a total of 4,992 model fits. We calculated the log pointwise predictive density (*lppd*) to score each model’s ability to predict the left-out data (Gelman et al. 2014). Thus, for training data *y*_train_ and held-out data *y*_hold_ at a given value of 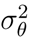 across all MCMC samples *s* = 1, 2, …, *S* and all hold outs of data from year *t* to year *T*, and letting *θ* represent all unknowns, *lppd* is

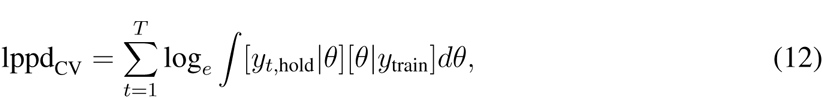

and computed as

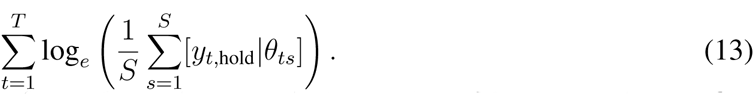

We chose the optimal prior variance for each species-statistical model combination as the one that produced the highest *lppd* and then fit each species-statistical model combination using the full data set for each species and the optimal prior variance. We calculated the *lppd* from posterior samples using the algorithm from Vehtari et al. (2016).

#### Population models

Using samples from the posterior distribution of the vital rate statistical models, it is straightforward to simulate the population models. We used an Integral Projection Model (IPM) to simulate populations based on individual-level data (Ellner and Rees 2006) and a quadrat-based version of an individually-based model (Quadrat-Based Model, QBM) to simulate populations based on quadrat-level data. We describe each in what follows.

##### Integral projection model

We use a stochastic IPM (Rees and Ellner 2009) to simulate our focal populations based on the vital rate regressions described above. In all simulations, we ignore the random year effects so that interannual variation is driven solely by climate. We fit the random year effects in the vital rate regressions to avoid over-attributing variation to climate covariates. Our IPM follows the specification of Chu and Adler (2015) where the population of species *j* is *n*(*u*_*j*_,*t*), giving the density of sized-*u* genets at time *t*. Genet size is on the natural log scale, so that *n*(*u*_*j*_,*t*)*du* is the number of genets whose area (on the arithmetic scale) is between *e*^*uj*^ and *e*^*uj+du*^. The function for any size *v* at time *t* + 1 is

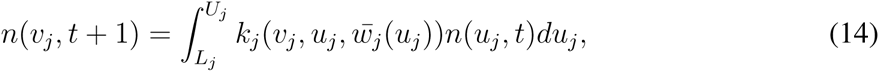

where 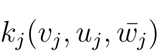, is the population kernel that describes all possible transitions from size *u* to *v* and 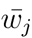 is a scalar representing the average intraspecific crowding experienced by a genet of size *u*_*j*_ and species *j*. The integral is evaluated over all possible sizes between predefined lower (*L*) and upper (*U*) size limits that extend beyond the range of observed genet sizes.

The IPM is spatially-implicit, thus, we cannot calculate neighborhood crowding for specific genets (*w*_*ij*_). Instead, we use an approximation 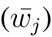 that captures the essential features of neighborhood interactions (Adler et al. 2010). This approximation relies on a ‘no-overlap’ rule for conspecific genets to approximate the overdispersion of large genets in space (Adler et al. 2010). The population kernel is defined as the joint contributions of survival (*S*), growth (G), and recruitment (*R*):

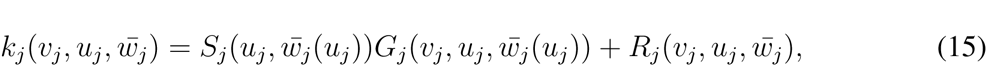

which means we are calculating growth (*G*) for individuals that survive (*S*) from time *t* to *t*+1 and adding in newly recruited (*R*) individuals of an average sized one-year-old genet for the focal species. Note the *S, G*, and *R* are incorporated in the IPM using the fitted vital rate regressions. Our statistical model for recruitment (*R*, described above) returns the number of new recruits produced per quadrat. Following previous work (Adler et al. 2012, Chu and Adler 2015), we assume that fecundity increases linearly with size 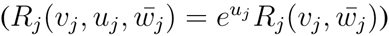 to incorporate the recruitment function in the spatially-implicit IPM.

We used random draws from the final 1,000 iterations from each of three MCMC chains for each vital rate regression to carry-through parameter uncertainty into our population models. At each time step, we drew the full parameter set (climate effects and density-dependence fixed effects) from a randomly selected MCMC iteration. Relatively unimportant climate covariates (those that broadly overlap 0) will have little effect on the mean of the simulation results, but can contribute to their variation. To retain temporal variation associated with random year effects, we used posterior estimates of the mean temporal effect and the standard deviation of that effect to generate a random year effect for unobserved years. That is, for some future year *T*, the intercept is *β*_0,*T*_ ~ Normal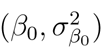 and the effect of size is *β*_*s*_,_*T*_ ~ Normal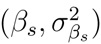.

##### Quadrat-based model

To simulate our quadrat-based model (QBM), we iterate the quadrat-level statistical model (Eqs. 9–10). We use the same approach for drawing parameter values as described for the IPM. After drawing the appropriate parameter set, we calculate the mean response (log cover at *t+*1 is *µ*_t+1_) according to Eq. 10. We make a random draw from a [0,1] truncated lognormal distribution with mean equal to *µ*_*t*+1_ from Eq. 10 and the variance estimate from the fitted model. We project the model forward by drawing a new parameter set (unique to climate year and MCMC iteration) at each timestep. Random year effects are included as described above for the IPM.

#### Model validation

To test each model’s ability to forecast population states, we made out-of-sample predictions using leave-one-year-out cross validation. For both levels of modeling and for models with and without climate covariates, we fit the vital rate models using observations from all years except one, and then used those fitted parameters in the population models to perform a one-step-ahead forecast for the year whose observations were withheld from model fitting. We made predictions for each observed quadrat in each focal year, initializing each simulation with cover in the quadrat the previous year. Because we were making quadrat-specific predictions, we incorporated the group random effect on the intercept for both models. We repeated this procedure for all 13 observation years, making 100 one-step-ahead forecasts for each quadrat-year combination with parameter uncertainty included via random draw from the MCMC chain as described above. As described above, year-specific parameters for left-out data were drawn from the posterior distribution of the mean intercept.

This cross-validation procedure allowed us to compare the accuracy and precision of the two modeling approaches (IPM versus QBM) with and without climate covariates. We first calculated the median predicted cover across the 100 simulations for each quadrat-year and then calculated forecast skill as the correlation (*ρ*) between forecasts and observations. We calculated forecast error as mean absolute error (MAE) between forecasts and observations. We compared *ρ* and MAE between model types and within model types between models with and without climate covariates using one-sided *t* tests with adjusted degrees of freedom following Wilcox (2009) and standard errors calculated using the HC4 estimator of Cribari-Neto (2004). Statistical tests for comparing correlations and error were conducted using algorithms from Ye et al. (2015).

#### Forecast horizons

An important feature of any forecasting model is the rate at which forecast skill declines as the time between an observation and a forecast increases. In particular, we are interested in the temporal distance at which forecast skill falls below a threshold: the so-called ecological forecast horizon (Petchey et al. 2015). To assess the forecast horizons of our models, we initiate the forecast model with the population state at some time *t* and make sequential forecasts of the population at times *t* + 1, *t* + 2, …,*t* + *T* where *T* is the maximum number of years between the initial year and the final year of our observations. For example, if we initialize the forecast model with percent cover in 1940, we are able to make five forecasts up to the year 1945. Forecast models are not re-initialized with observations between years. Thus, in our current example, the model forecast for percent cover in 1941 has a forecast horizon of one year, the forecast in 1942 has a forecast horizon of two years, and so on. We performed these simulations using mean parameter values for all model types (IPM with/without climate; QBM with/without climate) and all possible initial years. For a given forecast distance, we averaged the correlation between forecasts and observations. Note that our forecasts for the horizon analysis are all made using in-sample data because we used model fits from the full data set. Nonetheless, our simulations offer insight into the differences among model forecast horizons. We chose an arbitrary forecast accuracy of *ρ* =0.5 as our forecast proficiency threshold. the forecast horizon is the temporal distance at which forecast accuracy falls below *ρ* = 0.5. For basic research on forecasting, arbitrary proficiency thresholds suffice for comparative purposes (Petchey et al. 2015), and *ρ* = 0.5 represents the point at which about 25% of the variance in observations is explained by the predictions.

### Results

The IPM and QBM generated one-step-ahead forecasts of similar skill for out-of-sample observations, with an average correlation between predictions and observations (*ρ*) of 0.72 across all models and species (Fig. 2). Without climate covariates, the accuracy of forecasts from the IPM were not statistically greater than the accuracy of forecasts from the QBM (Fig. 2) and overall error was similar (mean absolute error; Fig. S1, Supporting Information). With climate covariates, the best out-of-sample predictive model (highest *lppd*) for each species and vital rate typically resulted from highly constrained priors on the climate effects (Fig. S2, Supporting Information). Thus, the posterior distributions of climate effects included in our models overlapped zero and generally were shrunk toward zero, though for some species-vital rate combinations, important effects (80% credible interval does not include zero) did emerge (Fig. 3).

**Figure 2.**
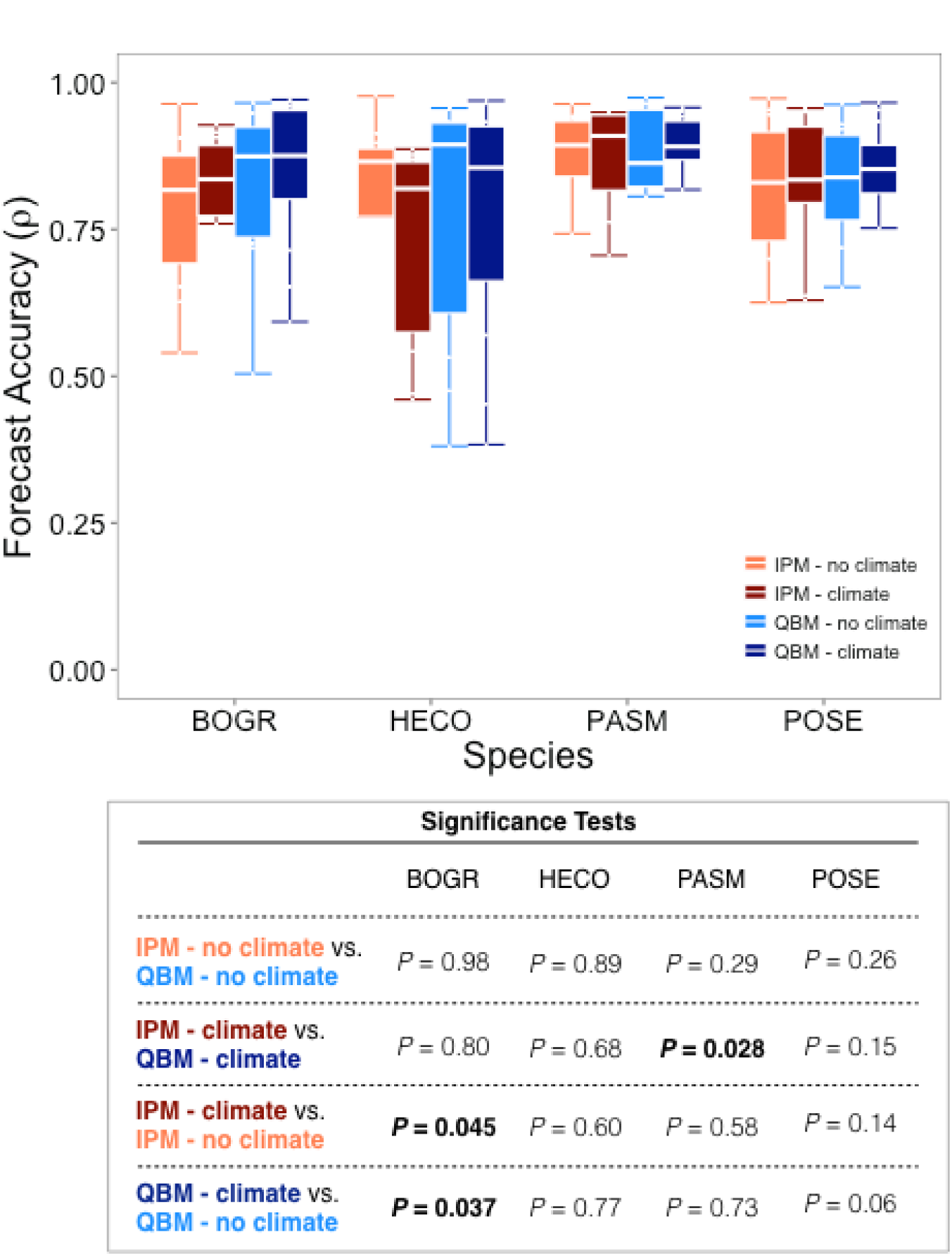
Comparisons of one-step-ahead, out-of-sample forecast accuracy between the IPM and QBM models with and without the inclusion of climate covariates. Boxplots show the distribution of *ρ* averaged over quadrats for each cross-validation year (i.e., 13 values of *ρ* for each species-model combination). For each comparison, *P*-values are from one-sided *t* tests designed to assess whether the first model in the comparison statement had higher accuracy than the second model in the comparison statement (see details in Table S22). Statistical tests relied on correlation values for each quadrat-year-species combination, after averaging over model reps for each combination. In no case did adding climate covariates decrease forecast accuracy (Table S21). Species codes are as in Fig. 1.

Despite the weak climate effects, including climate covariates did increase the accuracy of forecasts for two species: *B. gracilis* and *Poa secunda* (Fig. 2). However, only for *B. gracilis* were the skill increases statistically significant at *α =* 0.05 for the IPM (*t*_(279)_ = 1.70, *P* = 0.045) and the QBM (*t*_(279)_ = 1.80, *P* = 0.037). Similarly, forecast error decreased significantly with the inclusion of climate covariates for the *B. gracilis* IPM (*t*_(280)_ = −3.72, *P* = 0.029) and QBM (*t*_(280)_ = −3.34, *P <* 0.0001), and for the *Poa secunda* IPM (*t*_(196)_ = −1.90, *P <* 0.0001) and QBM (*t*_(196)_ = −2.47, *P* = 0.007) (Fig. S2, Supporting Information). In no case did including climate covariates significantly decrease forecast skill (Table S21), despite small changes in the mean skill (Fig. 2).

IPM forecasts were significantly more accurate than the QBM in only one case (Fig. 2): forecast accuracy of *P. smithii* percent cover from an IPM with climate covariates was greater than the accuracy from the QBM with climate covariates (*t*_(215)_ = 1.92, *P* = 0.028). However, adding climate covariates decreased the skill of both models, and the difference between the IPM and QBM emerges only because skill decreased less for the IPM than the QBM. Results from all pairwise statistical tests are shown in Table S22 of the Supporting Information.

With climate covariates included and using mean parameter values, the accuracy of both models’ forecasts declined as the distance between the last observation and the forecast increased, but they did so at similar rates (Fig. 4). The only exception is for *Poa secunda*, where QBM forecast accuracy remained steady as the temporal distance of the forecast increased, whereas IPM forecast accuracy declined (Fig. 4). The forecast horizons were short: forecast accuracy fell below *ρ* = 0.5 after one year for the IPM for most species, and after four years, at most, for the QBM (Fig. 4). Across the different temporal distances from the observation to the forecast, the IPM was never more accurate than the QBM (*P* > 0.05 for all one-sided *t*-tests, Table S23). Likewise, the QBM was rarely more accurate the IPM, the only exception being for *H. comata* at temporal distances of two (*t*_(115)_ = 2.39, *P* = 0.002) and three years (*t*_(98)_ = 2.04, *P* = 0.022) (Table S24). There were some cases where the QBM was more accurate than the IPM for *Poa secunda*, but neither model exceeded the forecast proficiency threshold by a large margin (Fig. 4, Table S24).

### Discussion

Our comparison between a traditional, demographic population model without environmental forcing (the IPM) and an equivalent model inspired by density-structured models (the QBM) showed that IPM forecasts of out-of-sample plant population states were no more accurate than forecasts from the QBM (Fig. 2; ‘no-climate’ bars). This result differed from our expectation that the IPM would out-perform the QBM, because of its mechanistic representation of the perennial life cycle. Our result also confirms theoretical (Freckleton et al. 2011) and empirical work (Taylor and Hastings 2004, Queenborough et al. 2011) showing that density-structured models can be useful surrogates for demographic models when the goal is to estimate or forecast population states over large spatial extents.

We also expected the inclusion of environmental forcing to reveal further differences between the models. Interannual variation in weather can affect vital rates in different ways (Dalgleish et al. 2011). Thus, estimates of climate effects on plant population growth may be biased or non-identifiable when the underlying statistical model is fit using population-level data that integrates over the potentially unique climate responses of individual vital rates. We found some evidence that the QBM failed to detect climate effects for three species (*B. gracilis*, *H. comata*, and *Poa secunda*), where important climate effects were identified in the individual vital rate models but not in the percent cover model (Fig. 3). For *H. comata*, adding climate covariates did not improve forecasts (Fig. 2), despite the significant climate effects in the vital rate regressions (Fig. 3).

**Figure 3.**
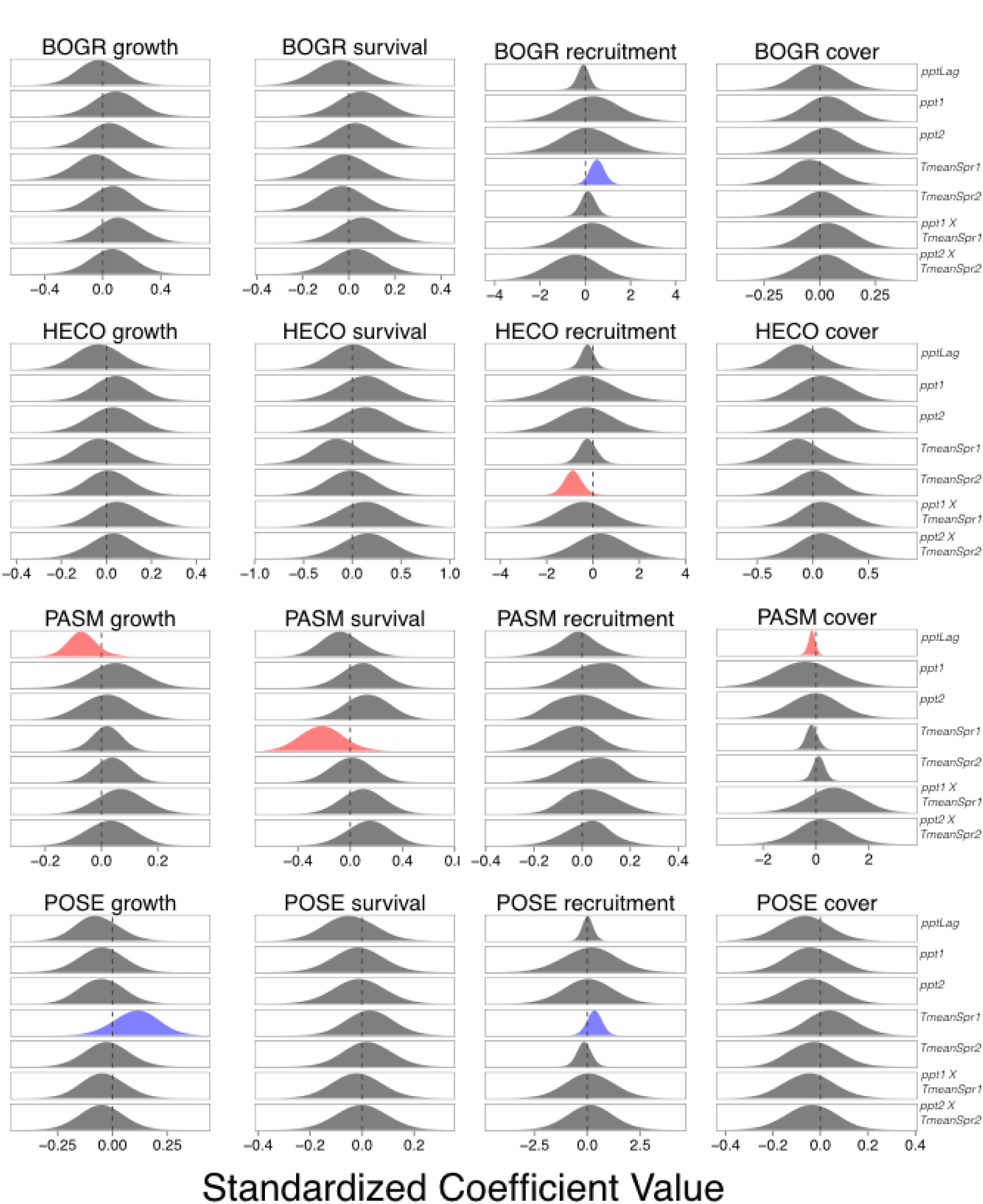
Posterior distributions of climate effects (*β*_*C*_) for each species and vital rate statistical model. Because our priors were constrained via ridge-regression, we highlight climate effects whose 80% credible intervals do not overlap zero (red for negative coefficients, blue for positive coefficients). Kernel bandwidths of posterior densities were adjusted by a factor of 4 for visual clarity. Species codes are as in Fig. 1. Climate covariate codes: *ppt*Lag = “water year” precipitation at *t*-2; *ppt1* = April through June precipitation at *t*-1; *ppt2* = April through June precipitation at *t*; *TmeanSpr1* = April through June temperature at *t*-1; *TmeanSpr2* = April through June temperature at *t*.

Furthermore, for the two species where including climate covariates increased forecast accuracy (*B. gracilis* and *Poa secunda*), forecast accuracy (Fig. 2) and error (Fig. S2) were equivalent between the IPM and QBM.

The higher accuracy of the IPM and QBM with climate covariates for *B. gracilis* and *Poa secunda* highlights the advantage of contemporary modeling and variable selection approaches such as ridge regression and LASSO over techniques that would exclude “non-significant” effects from final models. Ridge regression allows researchers to retain covariates whose effects may be difficult to identify in noisy data or short time series. This is especially important when forecasting the impacts of climate variability, where it is important to include the effects of forcing variables (e.g., temperature and precipitation) even if such effects are difficult to identify. Indeed, we failed to detect strong climate effects in the QBM for *B. gracilis* and *Poa secunda*, but including climate covariates still improved forecasting skill (Fig. 2). If a species is truly unresponsive to a given climate variable, statistical regularization techniques will shrink the mean and variance of a covariate estimate toward zero (Hooten and Hobbs 2015). Of course, regardless of what model selection approach is adopted, a critical step is identifying the appropriate candidate covariates, which we attempted to do based on our knowledge of this semi-arid plant community. However, the climate covariates we chose required aggregating daily weather data over discrete time periods. It is possible that we did not choose the optimal time periods over which to aggregate. New methods using functional linear models (or splines) may offer a data-driven approach for identifying the appropriate time periods over which to aggregate to produce a tractable set of candidate climate variables (Sims et al. 2007, Pol and Cockburn 2011, Teller et al. 2016).

We also expected IPM forecast accuracy to decline at a lower rate than the QBM as the time between the model initialization and the forecast increased. In principle, more mechanistic models should produce better predictions, especially under novel conditions (Evans 2012, Schindler and Hilborn 2015). In our case, the IPM explicitly models the influence of weather on recruitment and survival, effects that may be poorly represented in the QBM because recruitment and survival mainly affect small plants that contribute little to year-to-year changes in percent cover. Over longer time scales, the addition and subtraction of small plants could have large effects on population growth, so explicitly modeling these effects could contribute to a longer forecast horizon. However, we found no evidence that the forecast horizon for the IPM was greater than the QBM (Fig. 4). On the contrary, the QBM tended to have a slightly longer forecast horizon than the IPM for most species (Fig. 4). The QBM has fewer processes and parameters, which can reduce bias due to parameter uncertainty. As a result, the QBM may better capture near term dynamics when populations do not fluctuate widely, as in our case.

**Figure 4.**
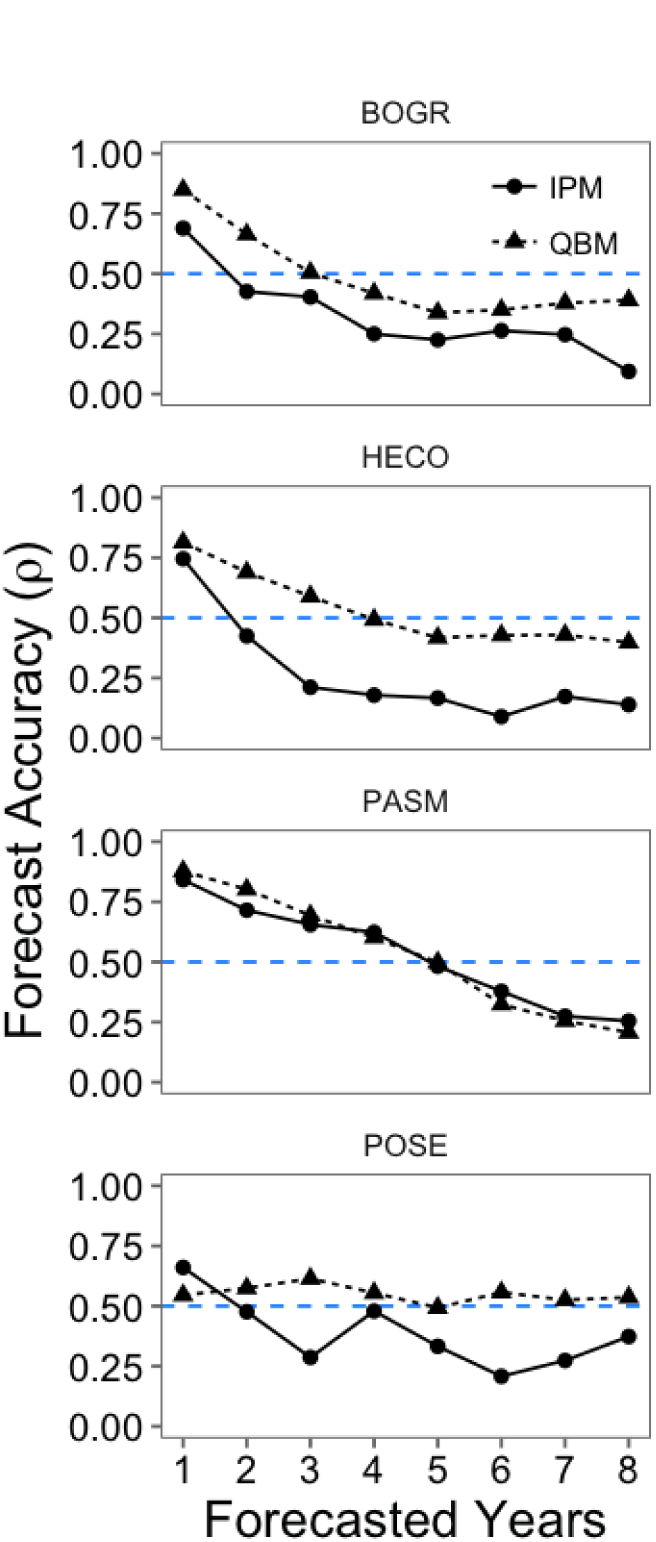
The forecast horizons for both models with climate covariates included and using mean parameter values. Points show the average accuracy (*ρ*, correlation between observations and predictions) across all forecasts at a given distance between the last observation and the forecast, where forecasts are made for in-sample data. We only examine the forecast accuracy of models with climate covariates included because in no case did including climate covariates significantly decrease accuracy (see Fig. 2). The dashed blue line indicates a forecast proficiency threshold of *ρ* = 0.5. Species codes are as in Fig. 1 and statistical comparisons between the IPM and QBM at each forecast distance are in Tables S23 and S24.

Our comparison of a model based on individual-level data with one based on percent cover data is not an exhaustive test. Understanding the reasons why the percent cover-based model matched the skill of a demographic model for our focal species may help us anticipate situations in which a percent-cover approach would fail. First, for none of our species did a climate covariate have a strong negative effect on one vital rate and a strong positive effect on a different vital rate (Fig. 3). As noted by Freckleton et al. (2011), complex age or stage structure can compromise predictions from models that aggregate over life-histories, and the same should be true when aggregating across vital rates with contrasting responses to climate drivers. Second, our particular recruitment model is already so aggregated – it averages across seed production, germination and establishment – that it may fail to detect important demographic responses to climate, putting our individual-based model and percent cover model on more equal footing. More finely resolved recruitment data might help our individual-based model outperform the population-level model. As advocated by Freckleton et al. (2011), knowledge of a species’ population ecology should guide the modeling approach. Third, our percent cover data are essentially error-free because we were able to aggregate indiviual plant areas to calculate percent cover. Percent cover data collected by typical sampling methods (e.g., Daubenmire frames) will include error that may affect population forecasts due to misspecifing the initial conditions and/or biasing model parameters (Queenborough et al. 2011). Percent cover models based on data containing more measurement error than ours might perform worse in comparison with individual-based models. One way to account for such error is to develop a sampling model that relates the observations (estimated percent cover in a plot) to the true state (percent cover derived from individual plant measurements in the same plot) (Hobbs and Hooten 2015).

Although our main goal was to compare individual-based and population-level modeling approaches relative to one another, it is worth reflecting on the absolute forecasting skill of our models. In particular, the forecast horizon of both models, defined as the time horizon at which the correlation between predictions and observations falls below *ρ* = 0.5, is less than five years for all species. Such short forecast horizons are not encouraging. Unfortunately, we have few ideas about how to improve population forecasts that have not already been proposed (Mouquet et al. 2015, Petchey et al. 2015). Longer time-series should improve our ability to detect exogenous drivers such as climate (Teller et al. 2016), and modeling larger spatial extents may reduce parameter uncertainty (Petchey et al. 2015). We may also have to shift our perspective from making explicit point forecasts to making moving average forecasts (Petchey et al. 2015). Whether the poor predictive ability of our models impacts the comparison of models based on individual vs. population-level data is an open question.

In conclusion, we found that models based on individual-level demographic data generally failed to generate more skillful population forecasts than models based on population-level data, even in models which included climate covariates. This finding runs counter to our expectations, but is consistent with recent theoretical (Freckleton et al. 2011) and empirical work (Queenborough et al. 2011). We conclude that models based on population-level data, rather than individual-level data, may be adequate for forecasting the states and dynamics of plant populations. This conclusion comes with the caveat that our analysis may be a weak test of the prediction that individual-level data is necessary for forecasting if different vital rates respond to climate in opposing ways, because climate effects were relatively unimportant in our vital rate regressions. Nonetheless, our results should encourage the use of easy-to-collect population-level data for forecasting the state of plant populations.

## Acknowledgments

This work was funded by the National Science Foundation through a Postdoctoral Research Fellowship in Biology to ATT (DBI-1400370), award MSB-1241856 to MBH, and a CAREER award to PBA (DEB-1054040). We thank the original mappers of the permanent quadrats in Montana and the digitizers in the Adler lab, without whom this work would not have been possible. Informal conversations with Stephen Ellner, Giles Hooker, Robin Snyder, and a series of meetings between the Adler and Weecology labs at USU sharpened our thinking. Brittany Teller provided comments that improved our manuscript. Compute, storage and other resources from the Division of Research Computing in the Office of Research and Graduate Studies at Utah State University are gratefully acknowledged. Any use of trade, firm, or product names is for descriptive purposes only and does not imply endorsement by the U.S. government. This research was supported by the Utah Agricultural Experiment Station, Utah State University, and approved as journal paper number 8917.

## Data Accessibility

The data used in this paper have been archived on Ecological Archives: http://esapubs.org/archive/ecol/E092/143/. All data and R code necessary to reproduce our work has been deposited on Figshare (http://doi.org/10.6084/m9.figshare.4007520) and is also available on GitHub (http://github.com/atredennick/MicroMesoForecast/releases/tag/v1.0).

## Author Contributions

ATT and PBA conceived the study; ATT, MBH, and PBA developed the statistical and modeling approach; ATT did the data analysis, with input from MBH and PBA;ATT wrote the first draft of the manuscript and all authors contributed to revisions.

## Footnotes

1 http://esapubs.org/archive/ecol/E092/143/

2 *Note to reviewers:* so that v1.0 will be associated with the published version of the manuscript, we have released v0.2 to be associated with this review version.

